# Revealing the genetic mechanisms underpinning the parasite-induced water-seeking behaviour of insects through RNA-seq

**DOI:** 10.64898/2025.12.18.694984

**Authors:** Upendra R. Bhattarai, Jean-François Doherty, Robert Poulin, Eddy Dowle, Neil J. Gemmell

## Abstract

The water-seeking behaviour in arthropods infected by mermithid nematodes or nematomorphs is a classic example of host manipulation, yet how these parasites induce the behavioural change is poorly known. We investigated the molecular mechanisms at the basis of this aberrant behaviour in European earwigs (*Forficula auricularia*) infected with the nematode *Mermis nigrescens*. We performed comparative RNA-seq analysis at different stages of infection and behaviour manipulation in both the host and the parasite. We detected a total of 12,876 and 9,722 expressed genes in the earwig and nematode, respectively. Differential gene expression (DGE) analysis showed 673 genes upregulated and 593 downregulated in earwigs, and 2,672 genes upregulated and 2,293 downregulated in nematodes across all comparisons between stages of infection and parasite sizes. Clustering analysis produced six and four clusters of differentially expressed genes in earwigs and nematodes, respectively, based on temporal patterns of expression across stages of infection. With overrepresented analysis using gene ontology terms, we found that the earwig shows increased signalling and sensory pathways, while the nematode ramps up transport and secretory genes during manipulation. We found shared GATA transcription factor motif enriched in the differentially over expressed genes during manipulation in both host and the parasite. Our study provides valuable insights into candidate pathways and genes in the host and the parasite, improving our understanding of the molecular regulation of water-seeking behaviour.

## Introduction

Animals continuously make behavioural choices impacting their fitness and reproductive success. Contrary to the assumption that behaviours are self-determined and naturally selected for the animal’s own fitness gain (McNamara et al., 2014; Travis & Reznick, 2018), instances exist where this is not the case (Hafer, 2016; Hughes & Libersat, 2019; Poulin, 2010). Parasites often manipulate host behaviour to confer advantages upon themselves (Heil, 2016; Poulin & Maure, 2015), a strategy that has independently evolved across all major parasite phylogenetic lineages (Brown, 1999). Such manipulated hosts essentially serve as extended phenotypes of the parasites (Bailey, 2012; Dawkins, 1982; Hughes et al., 2008).

The water-seeking behaviour in arthropod hosts infected by parasitic worms (nematodes and nematomorphs) illustrates this phenomenon (Ponton et al., 2011; Thomas et al., 2002). The evolutionary significance of this aberrant behaviour is particularly striking because of the apparent convergent evolution of this behaviour in parasites from two different phyla (Herbison et al., 2019a; Obayashi et al., 2021). Nematomorphs and mermithid nematodes must emerge from their terrestrial arthropod hosts into water or water-saturated substrates, to avoid desiccation, reproduce and thus complete their life cycle (Baker & Capinera, 1997).

Despite historical documentation dating back to over 140 years (McCook, 1884), the underlying mechanisms of parasite-induced water-seeking remain underexplored. Early molecular investigations highlighted significant parasite-host interactions during behavioural modification (Biron & Loxdale, 2013). Research by Thomas and colleagues (2003) identified neurotransmitter and neuromodulator alterations in the brain of the cricket *Nemobius sylvestris* infected by the nematomorph *Paragordius tricuspidatus.* Ponton and colleagues (2011) showed the infected *N. sylvestris* are positively phototactic. Biron and colleagues (2005, 2006) carried out proteomic studies in grasshoppers and crickets infected with the nematomorphs *Spinochordodes tellinii* and *Paragordius tricuspidatus* respectively, correlating proteins from the Wnt signalling pathway with behavioural change. Moreover, although dehydration superficially increases interactions with water, protein profiling does not support this as a driving mechanism by hairworms (Coates et al., 2025).

In contrast, much less is known about the mechanisms underpinning the similar behavioural manipulation induced by mermithid nematodes. Williams et al. (2004) found that the mermithid nematode *Thaumamermis zealandica* causes its amphipod host *Talorchestia quoyana* to burrow deeper into sand, down to water-saturated layers, following an increase in haemolymph osmolality after infection, suggesting some thirst-driven mechanism facilitating nematode emergence, yet this observation lacks a molecular explanation.

Transcriptomics offers insights into gene expression and physiological mechanisms dictating behaviour (Han et al., 2015). While high-throughput sequencing has advanced host-parasite interaction studies (Goodwin et al., 2016), the lack of comprehensive genomic data limits its application in non-model organism. Dual RNA-seq, which profiles gene expression of both host and parasite has been a popular approach (Choi et al., 2014; Ehret et al., 2017; Mukherjee et al., 2021), however this strategy often struggles with RNA quality disparities between host and the parasite when extracted together (O’Keeffe & Jones, 2019). To address these issues, we adjusted our protocols to process host and parasite RNA separately, facilitating accurate dissection and individual sequencing. We also use a developmental series of both parasite and the corresponding host. The use of a developmental time series across infections is crucial to differentiate between gene expression changes associated with infection and those associated with host manipulation. This research utilizes the European earwig (*Forficula auricularia*) and its parasitic mermithid nematode (*Mermis nigrescens*) to explore gene expression across different infection stages, aiming to elucidate the genetic mechanisms driving the observed water-seeking behaviour. Behavioural changes in earwigs occur primarily when mermithids have reached maturity (Herbison et al., 2019b). As many cases of host manipulation in insects involve the central nervous system (Hughes & Libersat, 2018), we hypothesize that substantial changes in the transcriptome should occur there. When their development within the earwig host is complete, these nematodes induce their host to seek water and enter it (Herbison et al., 2019b), allowing the worm to emerge where it needs to be to reproduce. Although some proteomic changes in earwig brains following infection have been identified (Herbison et al., 2019a), these only provide a partial window into the underlying mechanisms responsible for host manipulation, and our comprehensive genomic approach focusing on both the host and parasite across developmental stages aims to address this knowledge gap.

## Materials and methods

### Insect rearing

Adult earwigs (*F. auricularia*) were collected from the Dunedin Botanic Garden (−45° 51’ 27.59” S, 170° 31’ 15.56” E) and reared in a temperature-controlled room (Temperature: cycling from 15 to 12°C, day/night; Photoperiod of L:D 16:8) in the Department of Zoology, University of Otago, Dunedin, New Zealand. Earwigs were placed in a 20L glass container with a dirt bed, dahlia flower petals, and pieces of cardboard cuttings as cover. They were provided with food containing a 50:50 mixture of ground oats and commercial cat food *ad libitum*. Water was sprayed every second day to maintain moisture in the container. Earwigs were left at least a week to acclimatize before sampling.

### Sampling

Earwigs were sampled across four-time points (Uninfected, Early infection, Late infection, and During manipulation) and nematodes across three-time points (Early maturation, Late maturation, During manipulation) (Figure 1). Adult earwigs were snap frozen in liquid N2 and dissected in 1xPBS under a dissection microscope to confirm the state and stage of parasitism. Samples were handled quickly on dry ice and stored at -80 °C until further use. Categorization of the samples was done by measuring the largest nematode found in each earwig. Uninfected earwigs (UI) have no detectable nematode infection. Early infection earwig and early maturation nematode correspond to an infected earwig with nematode sized <2 mm. Late infection earwig and late maturation nematode correspond to an infected earwig with nematode sized 7-9 mm. During manipulation samples were obtained by observing earwigs inside their rearing container with a water pool (6 cm diameter and 0.5 – 1 cm depth) in darkness for 5-6 hours using a red light. Once a nematode was seen coming out of an earwig in water, both specimens were individually snap frozen in liquid N2. All samples were collected between January - July 2020, except for one sample in the early stage of infection which was collected the following year from the same field population following the same sampling procedure. One late-stage infection sample was infected with multiple nematodes and only the largest nematode from this sample was processed for sequencing. Earwigs were sexed based on their morphological characteristics, however, nematodes were not due to technical difficulties.

**Figure 1.**
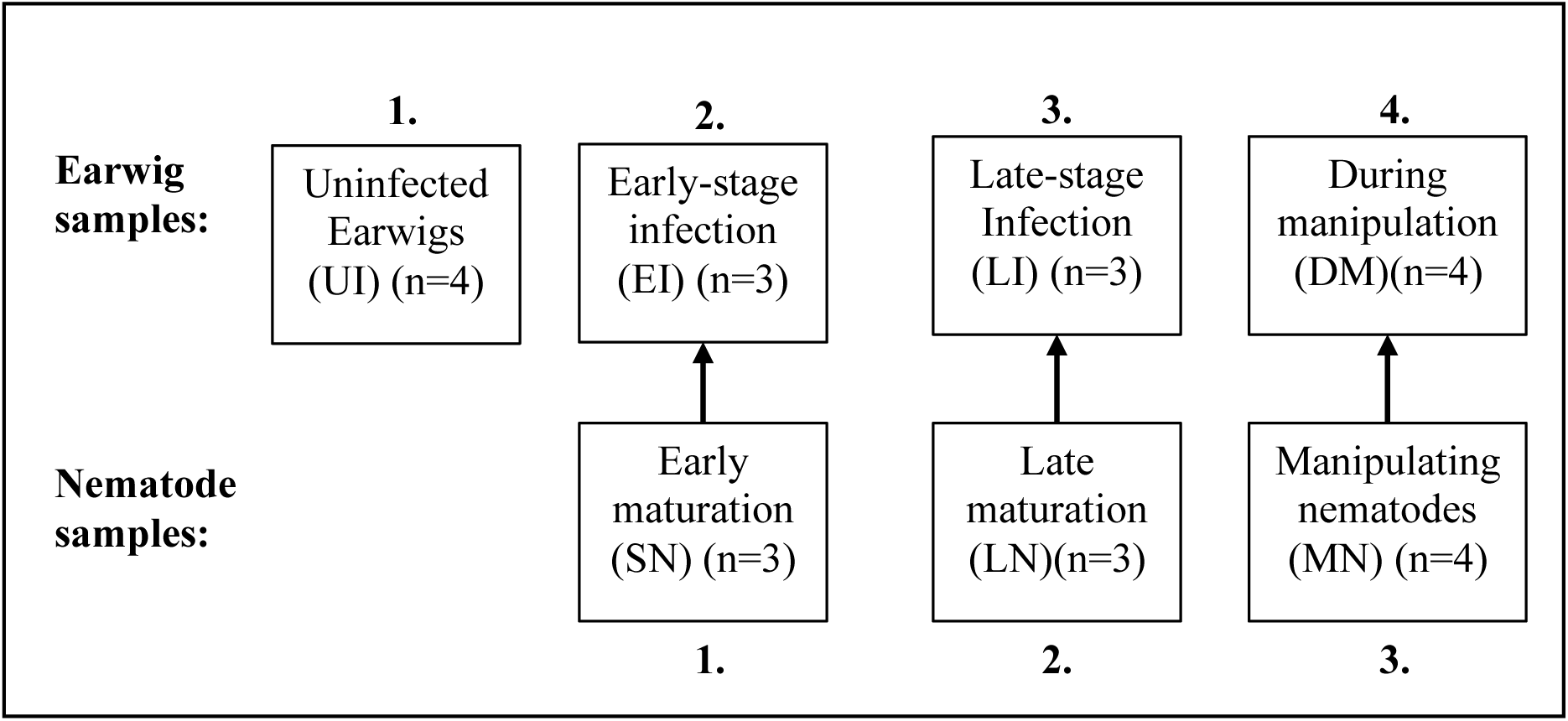
Sampling stages for RNA-seq studies of water seeking behaviour. Earwigs were sampled at four timepoints: Uninfected Earwigs (UI), Early-stage Infection (EI), Late-stage Infection (LI), and During Manipulation (DM) whereas, nematodes were sampled at three timepoints: Small Nematodes (SN) extracted from earwigs at early stage of infection, Large Nematodes (LN) extracted from earwigs at late stage of infection, and Manipulating Nematodes (MN) extracted from earwigs during manipulation.

### Nucleotide extraction

The total RNA extraction was carried out using Direct-zol RNA MicroPrep kit (Zymo Research, USA) with an on-filter DNAse treatment following the manufacturer’s instructions. Samples were ground in 300 μl 1x PBS buffer with blue sterile tips in a 2ml Eppendorf tube kept half dipped inside a mortar filled with liquid N2 to prepare for extraction. For earwigs, extractions were performed from their head (after removing the black eye pigments on the outer eye) to get the whole central nervous system, while for nematodes extractions were performed from their whole body. Quality and quantity of the extracted nucleotides were measured on a Qubit 2.0 Fluorometer (Life Technologies, USA) and Nanodrop. High quality RNA samples were stored at -80°C until use.

### RNA sequencing

RNA integrity was evaluated on a Fragment Analyzer (Advanced Analytical Technologies Inc., USA) and total RNA sequencing library was prepared with Zymo-Seq RiboFree Total RNA Library Kit (Zymo Research, USA). Prepared libraries were Quality checked on a MiSeq and sequenced across two lanes of Illumina HiSeq 2500 V2 Rapid Sequencing with 100bp paired end (PE) reads at the Otago Genomics Facility (OGF), University of Otago, Dunedin, New Zealand.

### Bioinformatics pipeline for RNA-Seq data

The RNA-seq reads were quality checked with FastQC and trimmed with Trimmomatic v.0.38 (Bolger et al., 2014) with the following parameters: ILLUMINACLIP:TruSeq3-PE-2.fa:2:30:10 LEADING:3 TRAILING:3 SLIDINGWINDOW:4:20 MINLEN:36. Reads were then mapped to their respective draft genomes (Bhattarai et al. 2022, 2024) using two pass mapping with the option --sjdbOverhang set to 99 in STAR v.2.7.9 (Dobin et al., 2013). RSEM v.1.3.3 (Li & Dewey, 2011) was used to generate read counts per gene with the rsem-calculate-expression command.

### Differential expression analysis and clustering

To understand how gene expression changed in both the earwig host and nematode parasite during infection a differential expression analysis was carried out in R (v.4.3.3) using DESeq2 package (Love et al., 2014). The sample variation for expression profiles was assessed using principal component analysis (PCA) with regularized log transformed data. For both earwig and nematode datasets, genes were retained if their expression count was >10 in at least three replicates. The deseq2 analysis was modelled to regress out the batch effects and account for both the sex and timepoints (design = ∼Batch + Sex + Timepoints) for earwig and timepoints as main effect for nematode dataset (design = ∼Batch + Timepoints). The pairwise differential expression was calculated between time points in both datasets using Wald test. Genes were considered differentially expressed when the corrected P value using the Benjamini and Hochberg method was below 0.05 and log2 fold-change was more or equal to 2. A non-duplicated list of all differentially expressed genes from each pairwise comparison was used for clustering analysis of their expression patterns across timepoints using the DEGReport package (v.1.38.5) (Pantano L, 2022) in R.

### Functional analysis

A over representation analysis (ORA) was performed with Gene Ontology (GO) terms for the genes in each cluster to identify common functional roles. GO annotation was extracted from the genome annotation (Bhattarai et al. 2022, 2024) and org.db database was created for each species using makeOrgPackage function from AnnotationForge (v.1.44.0) package. ORA was performed with enrichGO from clusterProfiler package (v.4.0) (Wu et al. 2021) with qvalue cutoff of 0.05 and multiple correction with Benjamini-Hochberg with all other default parameters unless otherwise stated. For genes in cluster 3, 5, and 6, default parameters did not yield enriched terms, so we used more lenient thresholds with multiple correction set to none and qvalue cutoff of 1. The universal gene set included all the genes in the DGE analysis.

### Motif analysis

Motif enrichment analyses were used to identify common transcription factor binding sites that may be involved in host manipulation. Analyses focused on the cluster of genes upregulated during manipulation compared to uninfected, early and late infection stages. Motif enrichment analysis was carried out using the findMotifs.pl tool from the HOMER package (v.5.1) (Heinz et al., 2010) in FASTA mode. For the genes of interest we took 500bp on both sides of the transcription start site (TSS) as foreground and the same genomic window from all the expressed genes were used as the background.

## Results

### Differential gene expression analysis

A total of 213.1 million 100bp paired end (PE) reads were generated from earwig samples and 155.7 million 100bp PE reads for nematode samples. Principal component analysis (PCA) showed that the samples clustered primarily by infection or development stage for both earwigs and nematodes along PC1 and PC2, while no clustering was observed based earwig sex (Figure 2A, B). In earwigs, 32% of sample variation was explained by PC1 and 17% by PC2, whereas in nematodes, 71% of variation was explained by PC1 and 12% by PC2. Nematode samples, particularly those from the manipulation stages (MN) exhibited clear separation from other stages while earwig samples formed three separate clusters, uninfected (UI), early and late infection stages (EI and LI) and during manipulation (DM).

**Figure 2.**
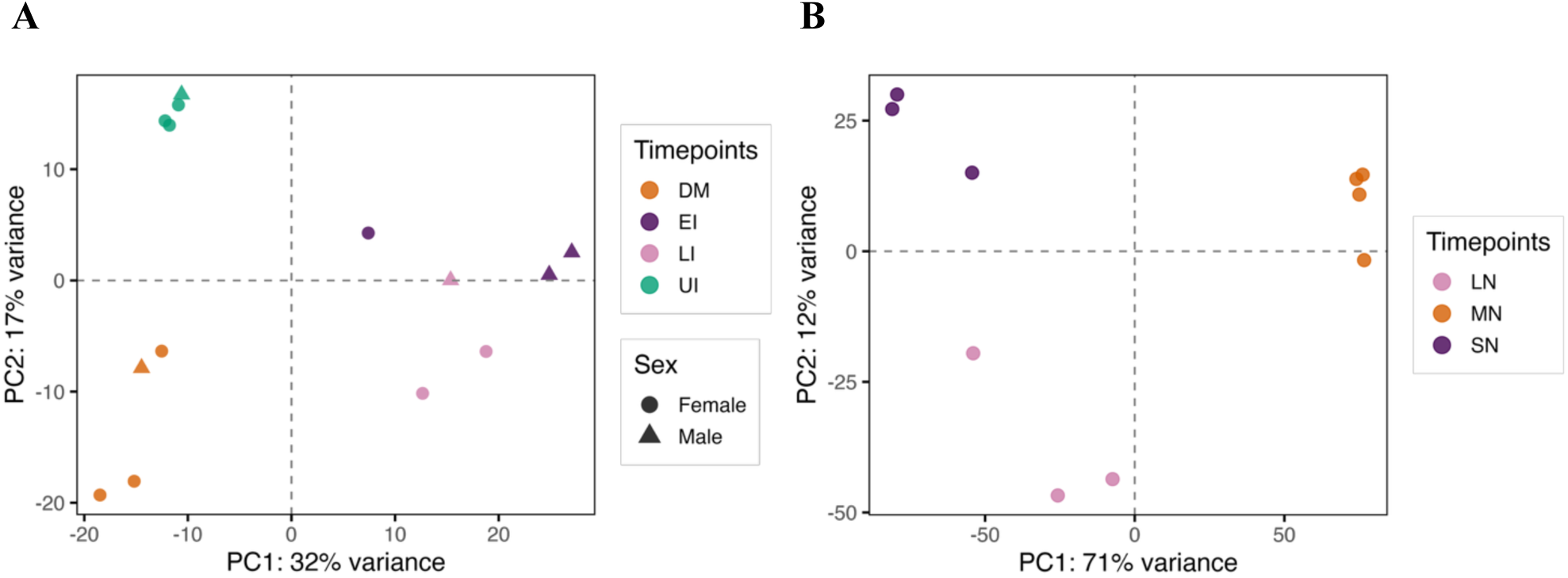
Principal Component Analysis (PCA) of the normalized RNA-seq data. A) Earwig samples showing groupings based on infection status with PC1 explaining 32% and PC2 17% of the variation. B) Nematode samples coloured by their progressive developmental stages, as determined by size and change induced in behaviour shows grouping based on the timepoints with PC1 explaining 71% and PC2 12% of the variation.

A total of 12,876 and 9,722 genes were detected in earwig and nematode datasets respectively. Among these 673 genes were significantly upregulated, and 593 genes downregulated in earwigs, and 2,672 genes were upregulated, and 2,293 genes downregulated in nematodes across all pairwise comparisons (Wald tests, FDR < 0.05 and log2FC ≥ 2). Heatmaps revealed clear clustering of samples by developmental stages in both the species (Figure 3A, B).

**Figure 3.**
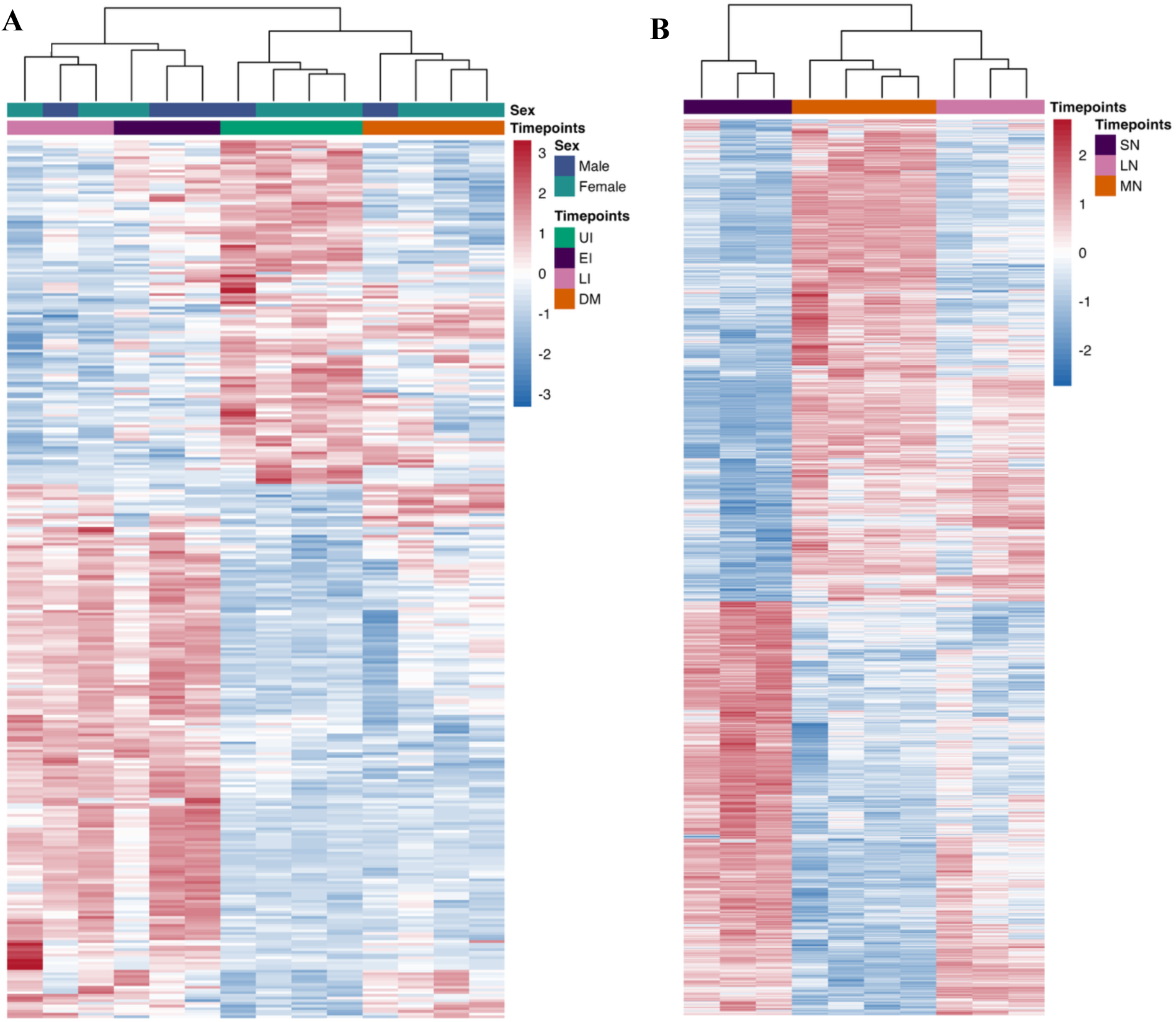
Heatmap showing normalized, rlog transformed read counts of top 500 differentially expressed genes (padj <0.05) for earwig samples (A) and nematode samples (B). Each row of the heatmap represents a gene differentially expressed across all samples colour coded according to Wald statistic test scores, where red indicates upregulation, and blue downregulation of a gene expression (gradient colour bar on the right). Colours above the heatmap indicate the sample categories as indicated by legends on the right.

Pairwise differential expression analysis in earwigs showed 34 upregulated and 93 downregulated genes between uninfected (UI) and Early infection (EI), whereas only 1 gene was upregulated and 5 downregulated between early (EI) and late infection (LI). Comparisons between late infection (LI) and during manipulation (DM) showed 187 upregulated and 46 downregulated genes. Similarly, 41 genes were up and 75 were downregulated between UI and LI, 162 up and 55 downregulated between UI and DM, and 256 up and 75 downregulated between EI and DM.

In the nematode dataset, 181 genes were upregulated, and 1,111 genes downregulated between small nematode (SN) and large nematode (LN); 609 up and 805 downregulated in large nematode (LN) and manipulating nematode (MN), and 761 up and 1,730 downregulated between SN and MN. Volcano plots illustrate the differences in magnitude and number of differentially expressed genes for each comparison (Supplementary Figure S1). In both the species, the changes of gene expression heightened during the manipulation stage.

### Clustering of DEGs based on expression patterns

The heatmaps of the DEGs revealed distinct gene groupings across the various timepoints in both the earwig and nematode datasets (Figure 3). To identify genes likely associated with host manipulations rather than those associated with infection or general nematode development DEGs were clustered based on their expression profiles across timepoints (Figure 4A, B).

**Figure 4:**
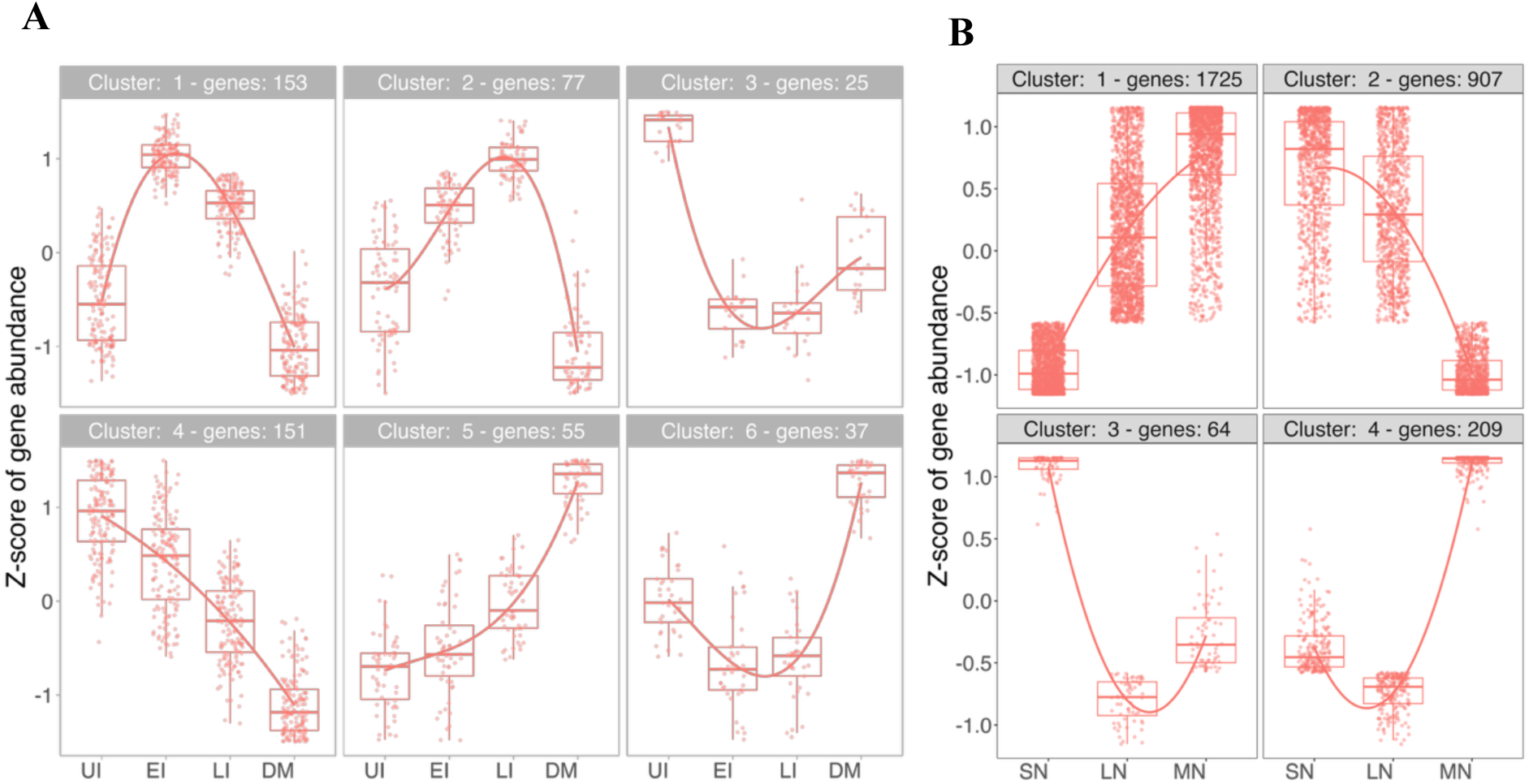
Cluster analysis of significantly expressed genes for earwig samples (A), nematode samples (B). Y- axis shows the Z-score of gene abundance, the values are centred to the mean and scaled to the standard deviation by each gene. X-axis shows different sampling stages. Each dot represents the expression of a gene; their relative expression values are summarized by box plots and whiskers at each sampling point. The curved lines represent the expressed trends. Number of genes in each cluster is given at the top of each cluster.

In the earwig dataset, six gene clusters with distinct patterns were identified. Cluster 1, 2 and 4 are characterized by reduced expression during the manipulation stage compared to earlier timepoints. In contrast, cluster 5 included genes showing a steady increase in expression from UI through EI, LI and in DM. Expression of the genes in clusters 3 reduced after infection to late infection and recovered slightly at DM, whereas, the genes in cluster 6 towered at DM after a slight decrease from UI to EI and LI. Cluster sizes ranges from 25 to 153 genes, with cluster 1 containing 153 genes, cluster 2 with 77, cluster 3 with 25, cluster 4 with 151, cluster 5 with 55, and cluster 6 with 37 genes. Of particular interest are cluster 2, which shows a pronounced drop in expression at manipulation stage, and cluster 6 which shows decreased expression following infection but rises during manipulation. These clusters may provide insights into the gene expression changes driving behavioural manipulation.

In the nematode dataset, DEG clustering yielded four groups with distinct temporal dynamics. Cluster 1 consists of genes that are consistently upregulated from the SN to the LN and MN stages, whereas cluster 2 shows the opposite pattern, with genes that are consistently downregulated across these stages. Cluster 3 and 4 are notable in that their genes exhibit decreased expression from SN to LN but then show a significant increase from LN to the manipulation stage (MN). The gene counts in these clusters are 1725 for cluster 1, 907 for cluster 2, 64 for cluster 3, and 209 for cluster 4. Since our primary focus is on gene changes occurring during the manipulation stage, clusters 3 and 4 with their marked upregulation at MN represent particularly promising groups for functional analysis.

Overall, these results highlight distinct temporal expression patterns in both host and parasite, providing focused candidate gene clusters for investigating the molecular mechanisms underlying behavioural manipulation (Figure 4A, B).

### Functional analysis of significantly expressed genes in the earwig host

A over representation analyses (ORA) of within clusters gene GO terms was used to identify key functions associated with each of the clusters. Earwig cluster 1 had significant enrichment in three gene ontology (GO) terms. The most prominent term was extracellular region (padj. 1.48e-08), which included genes *cda1*, *obst-e*, *chit1*, *cht10*, and ForAur_00000656. These same genes were also enriched for the chitin binding category (padj. 2.82e-07). Additionally, the carbohydrate metabolic process term was enriched (padj. 2.86e-03) with genes including *chit1*, *cht10*, *cda8*, *cda1*, *cht2*, *gusb*.

For cluster 2, extracellular region (padj. 0.004) and chitin binding (padj 0.012) incorporating *cda1*, *chit1*, *obst-e* genes were again enriched. Other terms enriched in this cluster include iron ion binding, heme binding, tetrapyrrole binding. There was notable overlap of genes across these terms, particularly *cyp15a1*, *cyp9e2*, *cyp4c1*, *agmo*, *obst-e* (Figure 5B). Cluster 4 presented 15 enriched GO terms, with many enriched terms overlapping with cluster 2 (Figure 5D).

**Figure 5.**
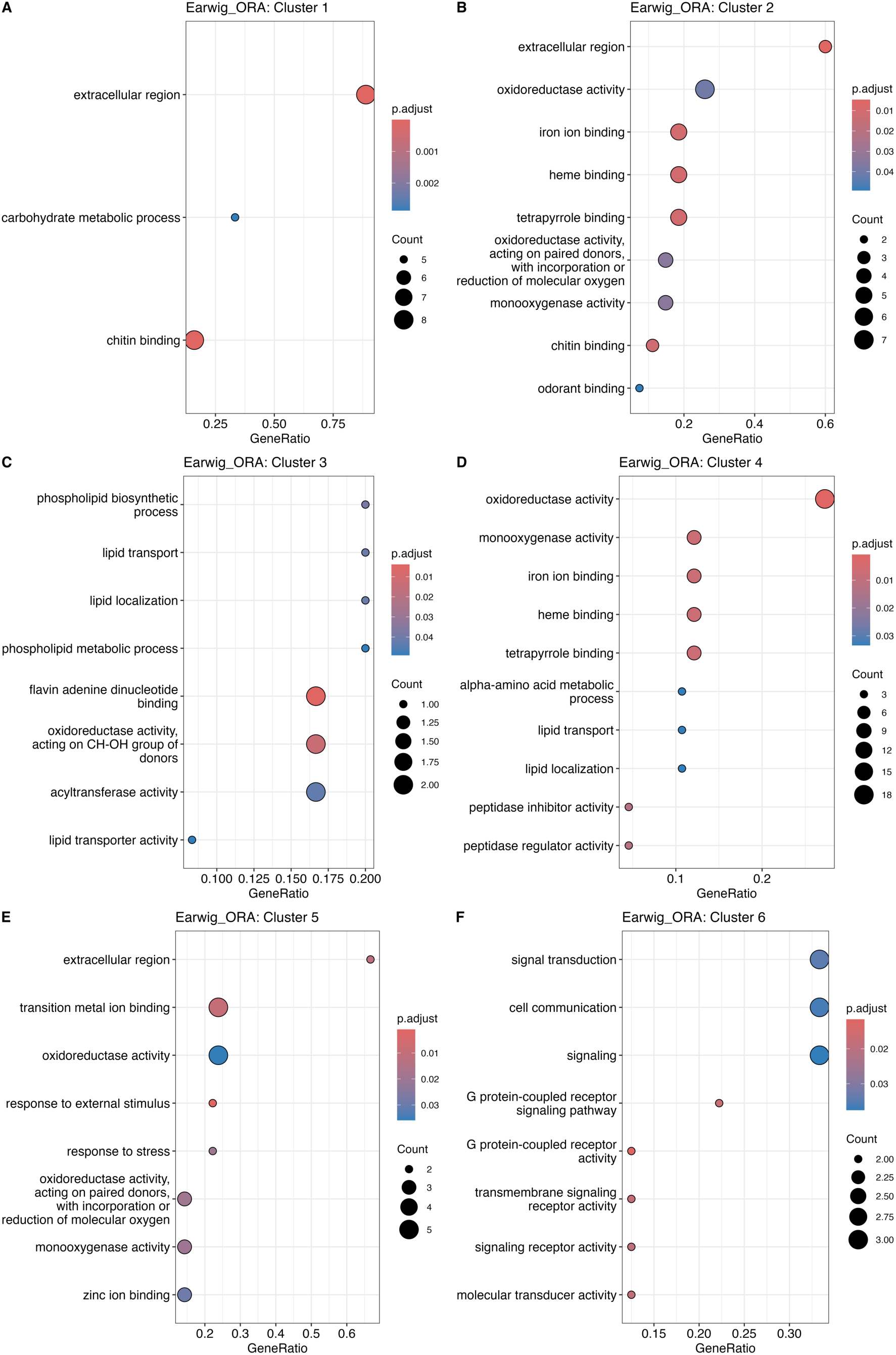
Dot plots showing top 10 enriched GO terms from the over representation analysis of genes in earwig clusters 1- 6. Adjusted p-values are shown by the colour gradient, red to blue for low to high, the size of the dot indicates the gene count of each term. X-axis represents gene ratio (scale varies between plots), and enriched terms are listed on the y axis.

Clusters exhibiting increased expression during the manipulation stage compared to EI and LI namely clusters 3 and 5 initially did not yield significantly enriched terms under the qvalue cutoff of <0.05 and Benjamini-Hochberg multiple correction. Using a more lenient threshold (pAdjustMethod = “none”, qvalueCutoff = 1), cluster 3 returned eight enriched GO terms, four of which belonged to the biological process category. Each featured a gene ratio of 1/5 with *vg2* associated with lipid transport and localization, and *hmgcs-1* involved in phospholipid biosynthetic and metabolic processes. The remaining four enriched terms were from the molecular function category, involving genes such as *gld*, *beta*, *vg2*, *nrf-6*, *oacyl*, *hmgcs-1* (Figure 5C). Cluster 5 also showed increased expression throughout infection and manipulation stages. Enriched GO terms included extracellular region, transition metal ion binding, oxidoreductase activity, response to external stimuli, response to stress, and others. (Figure 5D).

The most intriguing findings emerged from cluster 6, where gene expression was high during the manipulation stage after being downregulated during early and late infection (compared to uninfected stage). Most of the enriched terms in this cluster are related to signalling, particularly transmembrane signalling. These included signal transduction, cell communication, G protein-coupled receptor (GPCR) activity and pathway, transmembrane signalling receptor activity, and molecular transducer activity. Notably, just three genes *hcrtr2*, *adra1a, pde10a* were involved across all these enriched terms, and each exhibited higher expression during DM than at other time points (Figure 6).

**Figure 6.**
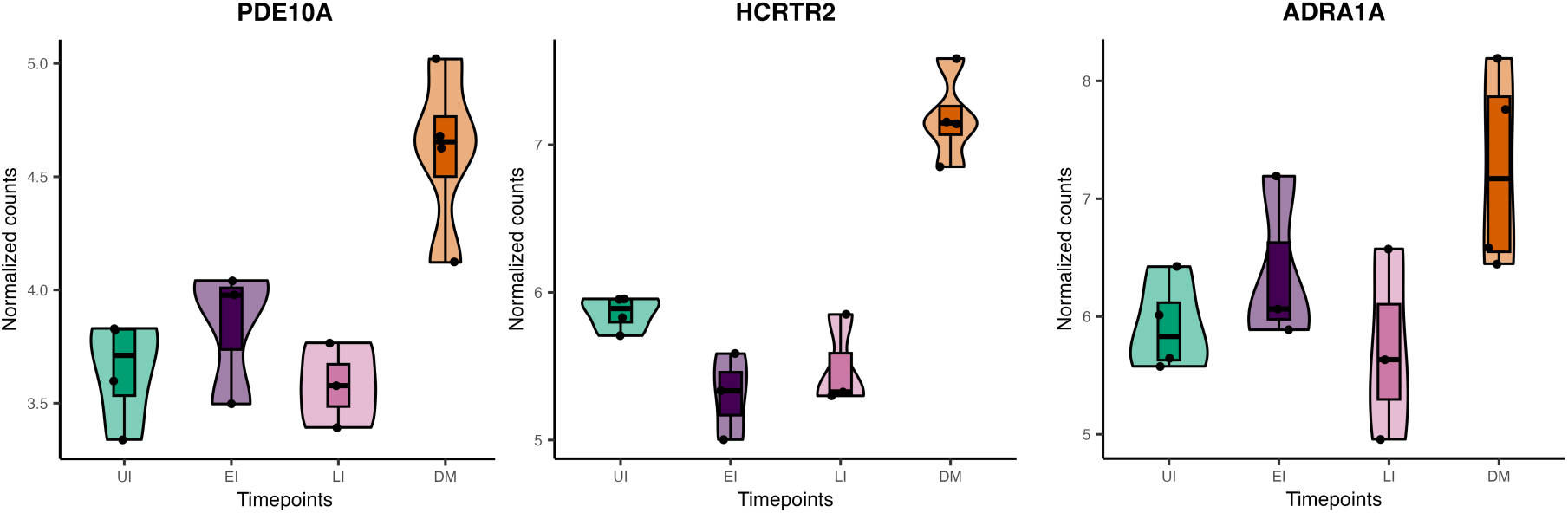
Expression of genes from enriched pathways in cluster 6 of the earwig dataset. A. *pde10A,* B. *hcrtr2,* C*. adra1a* gene expression. Y axis represents a log2 normalized gene counts and x axis represents the infection stages of the earwig. UI = uninfected, EI = in early infection stage, LI = in late infection stage, DM = During manipulation.

### Motif analysis of earwig genes putatively involved in host manipulation

Motif analyses were used within clusters with gene expression changes during manipulations to identify common gene regulatory regions. Enrichment of common motifs across candidate manipulation genes could suggest that the regulatory control of behavioural manipulations is linked to specific transcription factors. Motif analysis of genes in cluster 6 revealed nine known motifs (Table 1). Several of these motifs, including NFkB-p65, NFkB-p50, Fra2, TF3A are known regulators of immune responses and inflammation (Giridharan & Srinivasan, 2018, Sun, S.C, 2018, Patalano, D. S., Bass, P. F., Bass, J. I. F, 2023, Waldschmidt, R. et al. 1990). Among them, NFkB-p65 was the most enriched motif associated with 11 genes. Over representation analysis of GO terms for this gene set indicated enrichment for lipid metabolic process (padj. 0.03) and lipase activity (padj. 0.04), each represented by a single gene (*lip3* and *plbd2,* respectively) (Figure 7A).

**Table 1.**
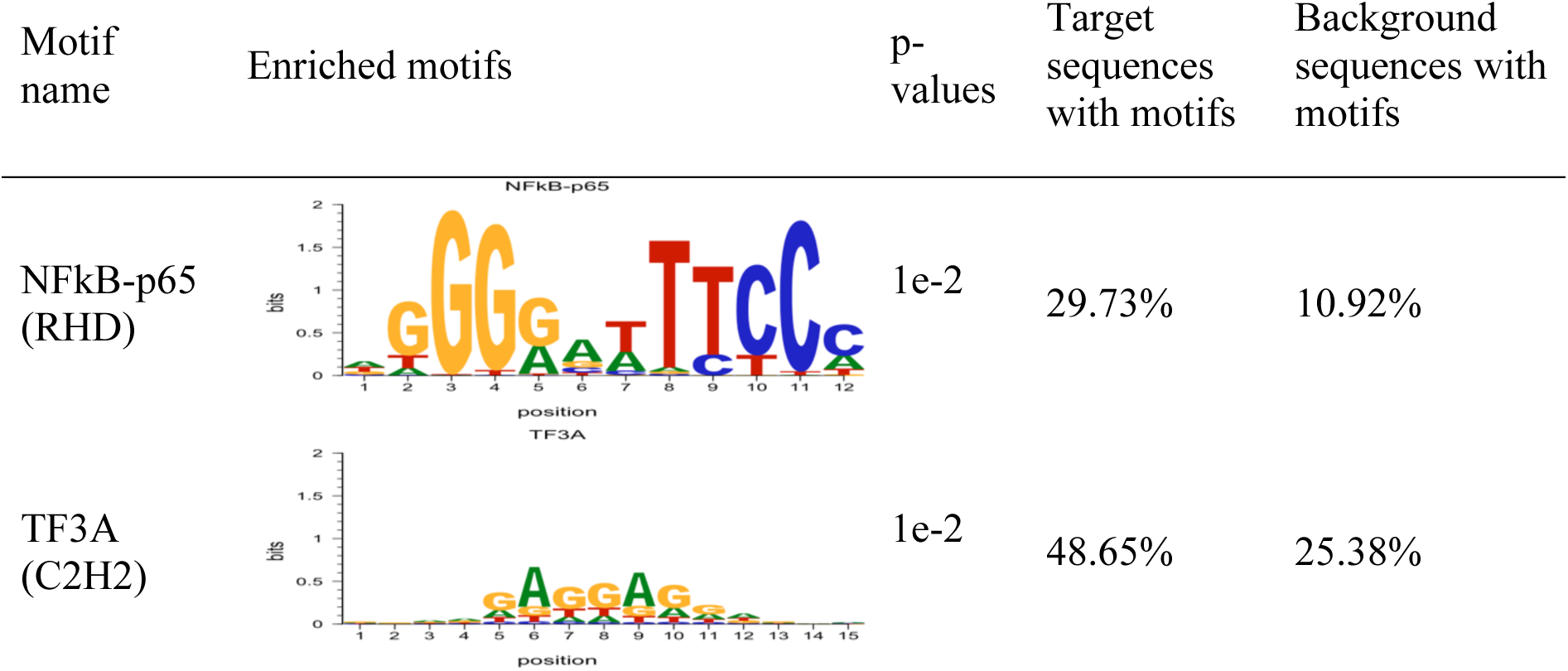
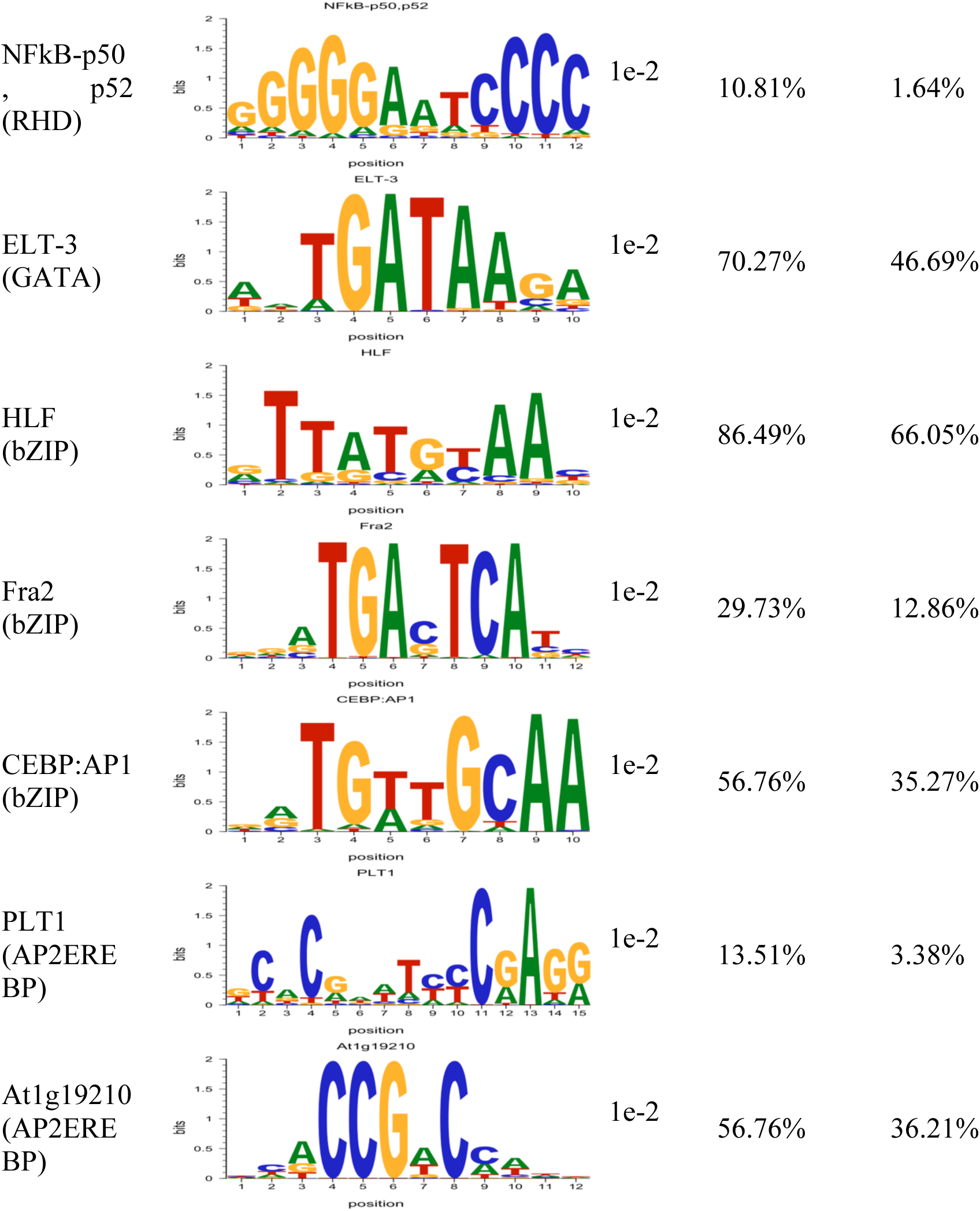
Known motif enrichment in genes from cluster 6 of the earwig dataset.

**Figure: 7.**
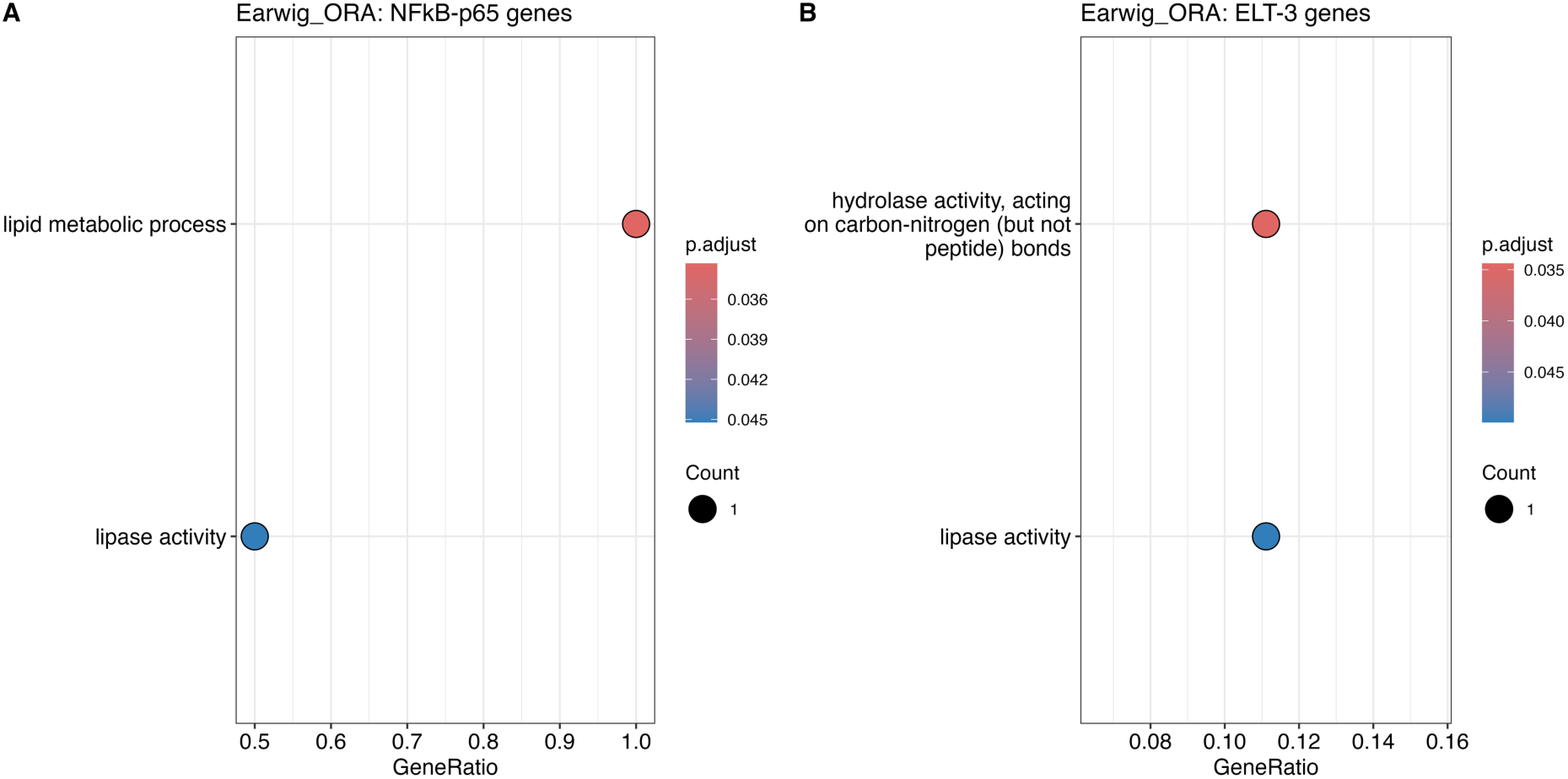
Dot plots showing enriched GO terms from the over representation analysis of genes potentially linked to manipulation from two of the overrepresented motifs. A) Genes with NFkB-p65 motif enrichment, B) Genes with ELT-3 motif enrichment. Adjusted p-values are shown by the colour gradient, red to blue for low to high, the size of the dot indicates the gene count of each term. X-axis represents gene ratio, and enriched terms are listed on the y axis.

The GATA transcription factor ELT3, a known regulator in age regulation in worms (Budovskaya et al. 2008) is surprisingly one of the enriched motifs in cluster 6 genes in the earwig. ORA using a lenient threshold has earlier identified two enriched GO terms: hydrolase activity, acting on carbon-nitrogen bonds (padj. 0.03) and lipase activity (0.04), each represented by one gene (*pgrp* and *plbd2,* respectively) (Figure 7B).

### Functional analysis of significantly expressed genes in the nematode

Clusters 1 and 2 in the nematode dataset showed distinct expression trends across time, with cluster 1 genes exhibiting progressively increased expression and cluster 2 genes showing a decrease from SN to MN. ORA identified 49 significantly enriched GO terms for cluster 1 and 23 for cluster 2. For cluster 1, top terms highlighted robust involvement in signalling and cellular communication, including pathways related to GPCR signalling and transmembrane transport (padj. 9.8e-06), cell communication (padj. 1.52e-05), signalling (padj. 1.6e-05), and signal transduction (padj. 4.33e-05). Additional terms pointed to heightened cellular responses to external stimuli (padj. 2.41e-03). Together, these results suggest a pronounced increase in transmembrane signalling activity in this group.

The most prominent features of cluster 2 were terms associated with biosynthetic and metabolic processes. The most significant terms included translation (padj. 4.69e-19), and the following terms were peptide and amide biosynthetic and metabolic process. This pattern indicates widespread downregulation of biosynthetic and metabolic activity in cluster 2 (Figure 8).

**Figure 8.**
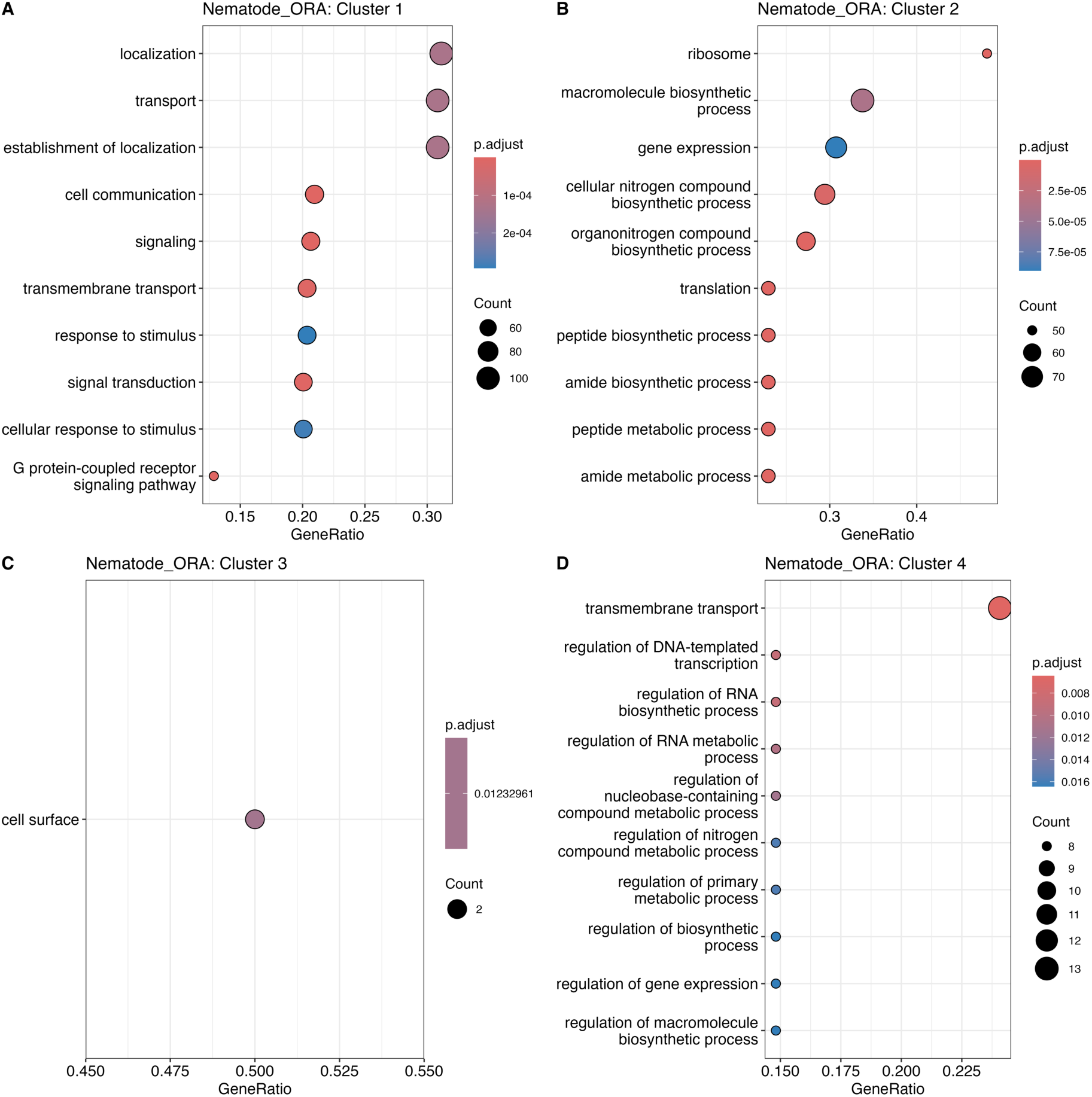
Dot plots showing the top 10 enriched GO terms from the over representation analysis of genes in nematode clusters 1- 4. Adjusted p-values are shown by the colour gradient, red to blue for low to high, the size of the dot indicates the gene count of each term. X-axis represents gene ratio, and enriched terms are listed on the y axis.

Cluster 3, characterized by a modest increase in expression at the manipulation stage (MN) following a decrease from SN to LN, yielded a single significantly overrepresented term, cell surface (padj. 0.01) including the genes *ttr-52* and *ttr-2*. In cluster 4, where expression strongly increased at MN after a slight decline from SN to LN, standard threshold did not yield significant enrichment. However, by relaxing the threshold as before, 25 enriched terms were identified. The most notable among these were transmembrane transport (padj. 0.005), regulation of DNA templated transcription and RNA biosynthetic process (padj. 0.014), and various metabolic and gene regulatory processes.

Within the transmembrane transport category, the majority of enriched genes encoded solute carrier proteins, such as *slc15a2*, *slc2a3*, *slc17a5*, *slc18a2*, *slc39a10*, as well as other transport related proteins like *abcc3*, *oatp74d*, *pept-3*, *trpc2*, *kha1*, *orct*, and *cac*. Many top enriched terms also consistently featured as set of regulatory genes, including *lrrfip2*, *meis3p2*, *ehf*, *foxp1*, *foxo2*, *st18*, *sgf1*, *eip74ef* (Figure 8).

### Motif analysis of nematode genes putatively involved in host manipualtion

Motif analysis of nematode cluster 4 genes identified six enriched known motifs: ABF1, GATA (Zf) IR3, AGL95 (ND), NFE2L2 (bZIP), Bach1 (bZIP), PHA-4 (Forkhead) (Table 2). Among these, GATA-IR3, a zinc finger DNA binding motif, though evolutionary distant shares fundamental structural similarities with ELT-3, the GATA zinc finger motif enriched in earwig cluster 6 genes. ORA using this motif revealed seven enriched GO terms at a lenient threshold as earlier (Figure 9), each represented by a single gene. These included UDP-glycosyltransferase activity, extracellular ligand-gated monoatomic ion channel activity, and oligosaccharide metabolic process containing gene *tre1*, ligand-gated monoatomic ion channel activity, ligand-gated channel activity and hydrolase activity acting on glycosyl bonds containing gene *exp-1*, and hydrolase activity, hydrolysing O-glycosyl-compounds containing gene *ugt3*.

**Figure 9.**
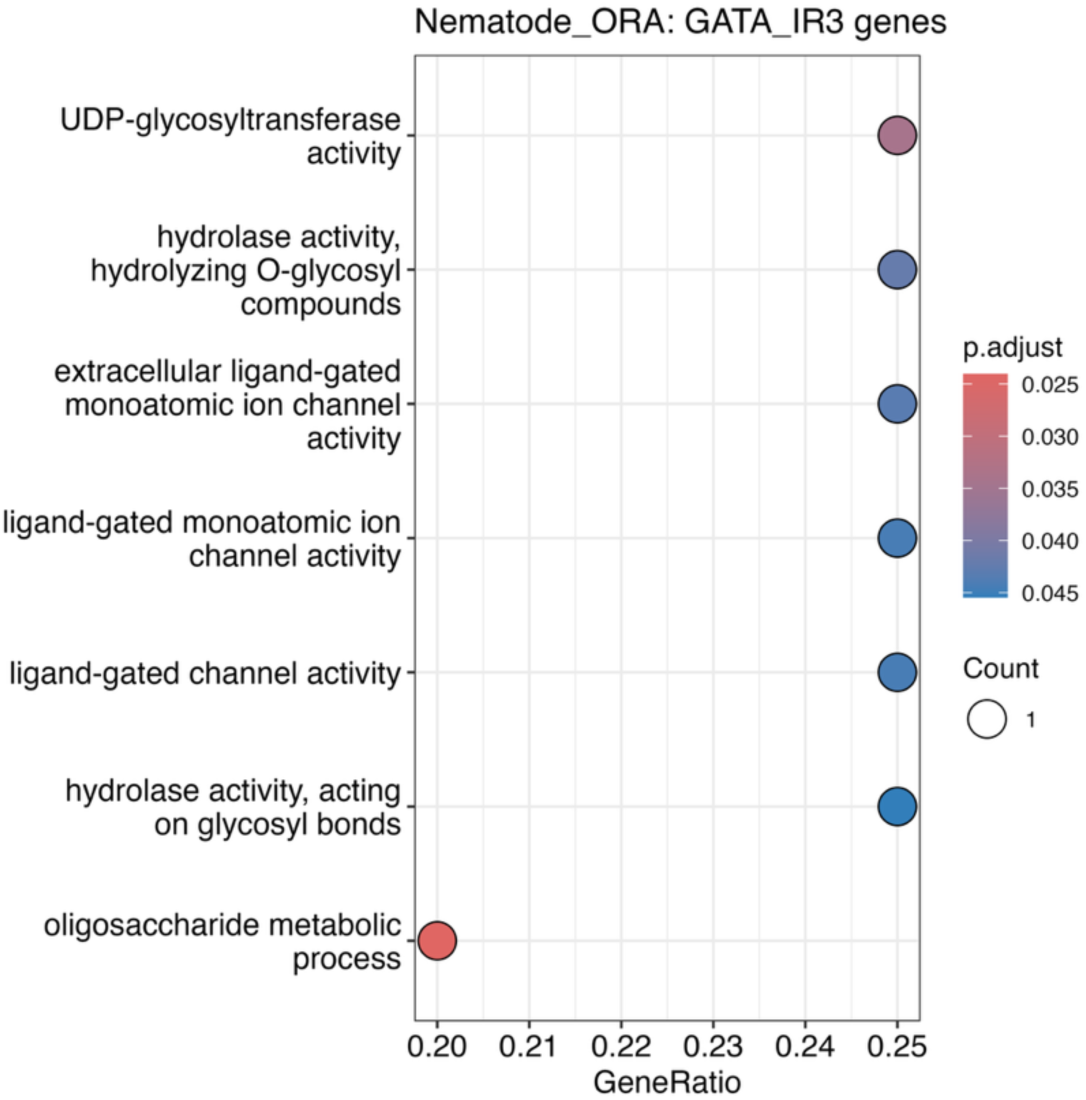
Dot plots showing enriched GO terms from the over representation analysis of genes with GATA_IR3 motif enrichment. Adjusted p-values are shown by the colour gradient, red to blue for low to high, the size of the dot indicates the gene count of each term. X-axis represents gene ratio, and enriched terms are listed on the y axis.

**Table 2.**
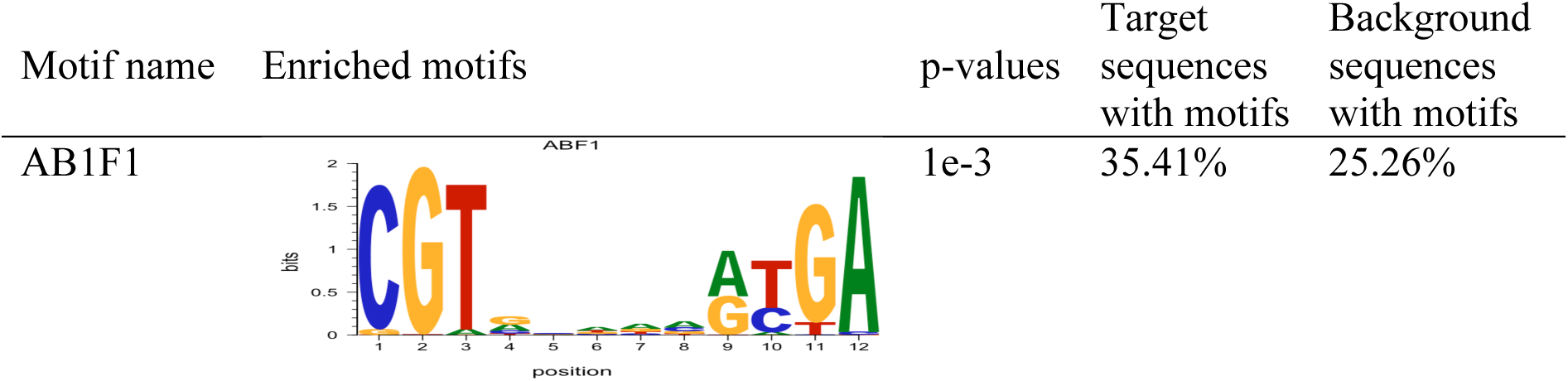
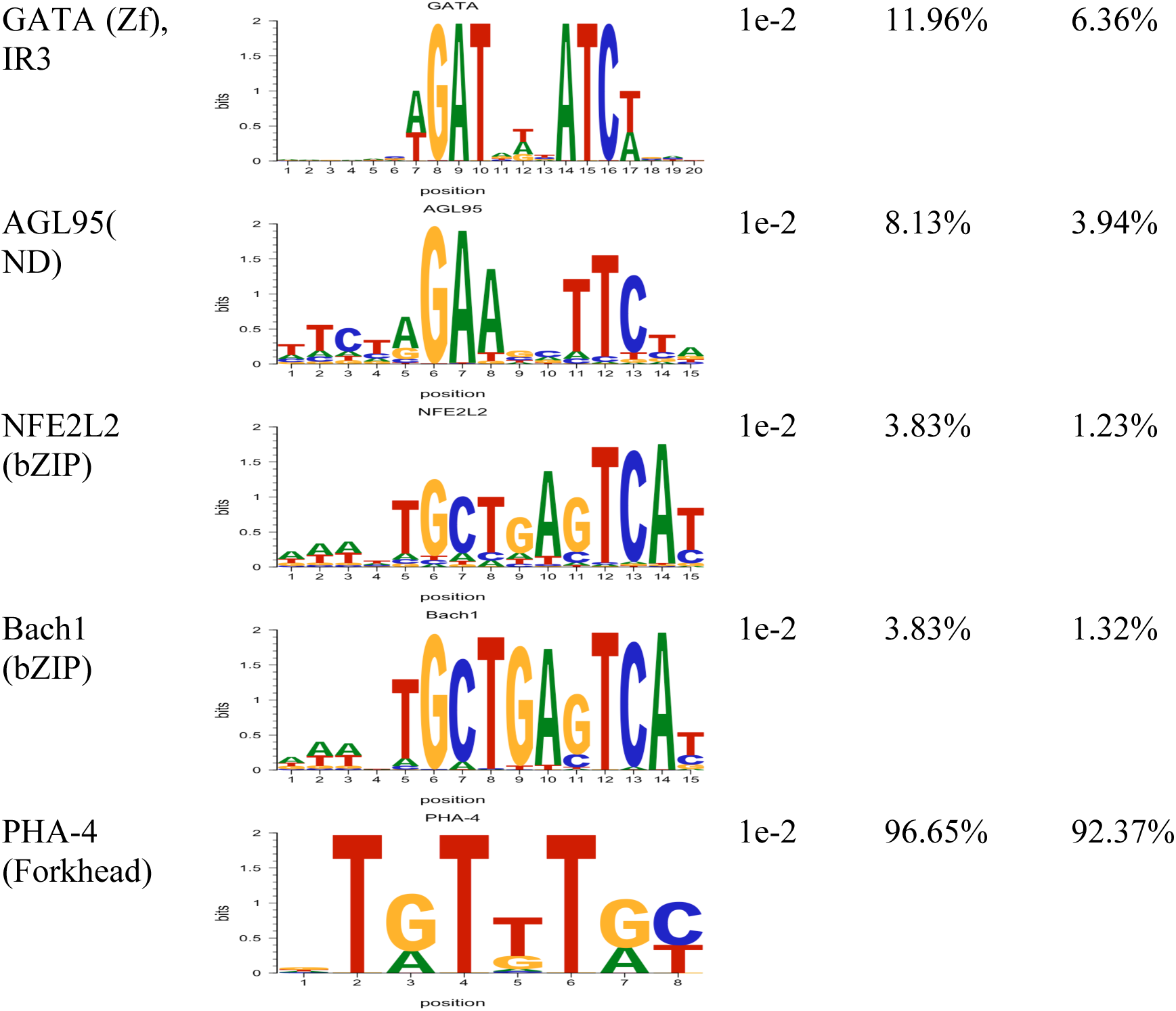
Known motif enrichment in genes from cluster 4 of the nematode dataset.

## Discussion

We hypothesized that the nematode manipulates earwig behaviour by interfering with the host’s central nervous system (CNS), thereby altering information processing and decision making by the host. In earwigs, early versus late infection produced relatively few differentially expressed genes. This limited change may reflect the nematode’s growth inside the abdomen having only subtle effects on the CNS until the parasite reaches the manipulation stage. In contrast, nematode samples represented distinct developmental stages, which likely explains the larger number of differentially expressed genes observed between stages. This pattern supports the notion that nematodes remain “under the radar” until they are fully developed and ready to emerge, an adaptive strategy critical for completing their lifecycle.

Focusing on the genes differentially expressed during manipulation, functional analysis revealed enriched signalling processes in both species. In earwigs, genes involved in transmembrane signalling were upregulated in the CNS, while nematodes showed increased activity of genes linked to carrier functions, molecular transport, and regulatory process. These complementary changes suggest a coordinated interaction in which parasite driven transport and regulatory mechanisms may modulate host signalling, ultimately producing the behavioural shifts observed.

Several host genes emerged as particularly noteworthy. In cluster 6 of the earwig data, *hctr2*, *adra1a*, and *pde10a* were strongly associated with enriched GO terms. *Pde10a,* which encodes phosphodiesterase 10A, is a cyclic nucleotide phosphodiesterase involved in signal transduction and dopamine receptor-associated second messenger pathway (Day et al. 2005, Bonate et al. 2022). Its role in cGMP hydrolysis is well established in both vertebrates and invertebrates (Day et al., 2005; Francis et al., 2000), and loss of *pde10a* expression has been linked to reduced locomotion and cognitive defects in mice (Hebb et al., 2008; Menniti et al., 2021). Notably, earwigs exhibit increased walking during the manipulative stage (personal observation), which may increase their chances of encountering water. This behavioural correlation highlights the *pde10a* gene as an interesting candidate gene for future investigation. The other genes, *adra1a*, encoding the adrenoceptor alpha 1A protein, and *hcrtr2* encoding hypocretin receptor 2 proteins are involved in G-protein coupled receptors (GPCRs), a family of transmembrane proteins involved in diverse physiological and regulatory functions, including immunity and sensory perception and immune responses in insects (Scott et al., 2011, Liu et al. 2021). Their expression further underscores the role of CNS signalling pathways in host manipulation.

On the parasite side, cluster 4 nematode genes were enriched for transmembrane transport and transcriptional and translational regulation. These processes are consistent with the secretion of molecules that could influence earwig signalling. Motif analysis revealed enrichment of conserved regulatory elements, including the zinc finger GATA motif in both species, and NFkB-p65 as the most enriched in earwig. The lipid metabolism related pathways were enriched in the earwig genes with those motifs. Lipid metabolism is tightly linked to neural function and cognitive processes (Vallochi et al. 2018). One of the genes enriching lipase activities, lip3 is a regulator of metabolic responses to starvation and aging (Hanschke et al. 2022). The host’s candidate genes with ELT-3 motifs enriching the functional GO terms are *pgrp* and *plbd*2. The *pgrp* gene encodes peptidoglycan recognition proteins (PGRPs); these are highly conserved pattern recognition proteins and are known to activate immune and neurobehavioral pathways (Lanz-Mendoza & Contreras-Garduno 2022, Li et al. 2025).

In nematodes, GATA-IR3 motif enriched genes includes *tre1*, which is a GPCR, essential for germ cell migration in *Drosophila* (Kunwar et al. 2003); similarly, exp-1 is important for nutrient uptake in parasitic worms (Mesen-Ramirez et al. 2019). The *ugt3* (UDP-glucuronosyltransferase) is involved in detoxification and longevity (McElwee et al. 2004) and may also contribute to the nematode’s ability to survive within the host and influence host signalling.

Taken together, our findings reveal that both earwigs and nematodes undergo coordinated transcriptional changes during the manipulation stage. Host genes linked to CNS signalling and parasite genes involved in transport, secretion, and regulation converge on shared functional pathways. The enrichment of conserved cis-regulatory motifs across both species suggests common molecular switches that may mediate cross-species communication. This interplay between host signalling and parasite regulation provides a molecular framework for understanding the remarkable behavioural manipulation observed in this system and highlights candidate genes for future functional validation.

## Acknowledgements

This study was funded by Royal Society Te Aparangi Marsden Fund grant (UOO1613) and the Departmental PhD scholarship from the Department of Anatomy, University of Otago. We thank Joanne Gillum and Sara Ferreira for their technical assistance, support staffs at the Dunedin Botanical Garden for their help in collecting earwigs. We acknowledge the use of New Zealand eScience Infrastructure (NeSI) high-performance computing facilities.

## Author Contributions

Conceptualization: N.J.G., R.P., E.D.; Funding: N.J.G., R.P.; Sampling: U.R.B., J.F.D.; Lab work and sequencing: U.R.B.; Bioinformatics and statistical analysis: U.R.B., E.D.; Writing: U.R.B., E.D., N.J.G. Revisions: all authors.

## Data Availability Statement

Raw sequence reads for both the host, and the parasites are deposited in the SRA repository under the BioProject PRJNA1364760.

