## Supplementary Figure S1 for "Revealing the genetic mechanisms underpinning the parasite-induced water-seeking behaviour of insects through RNA-seq"

### Supplementary materials

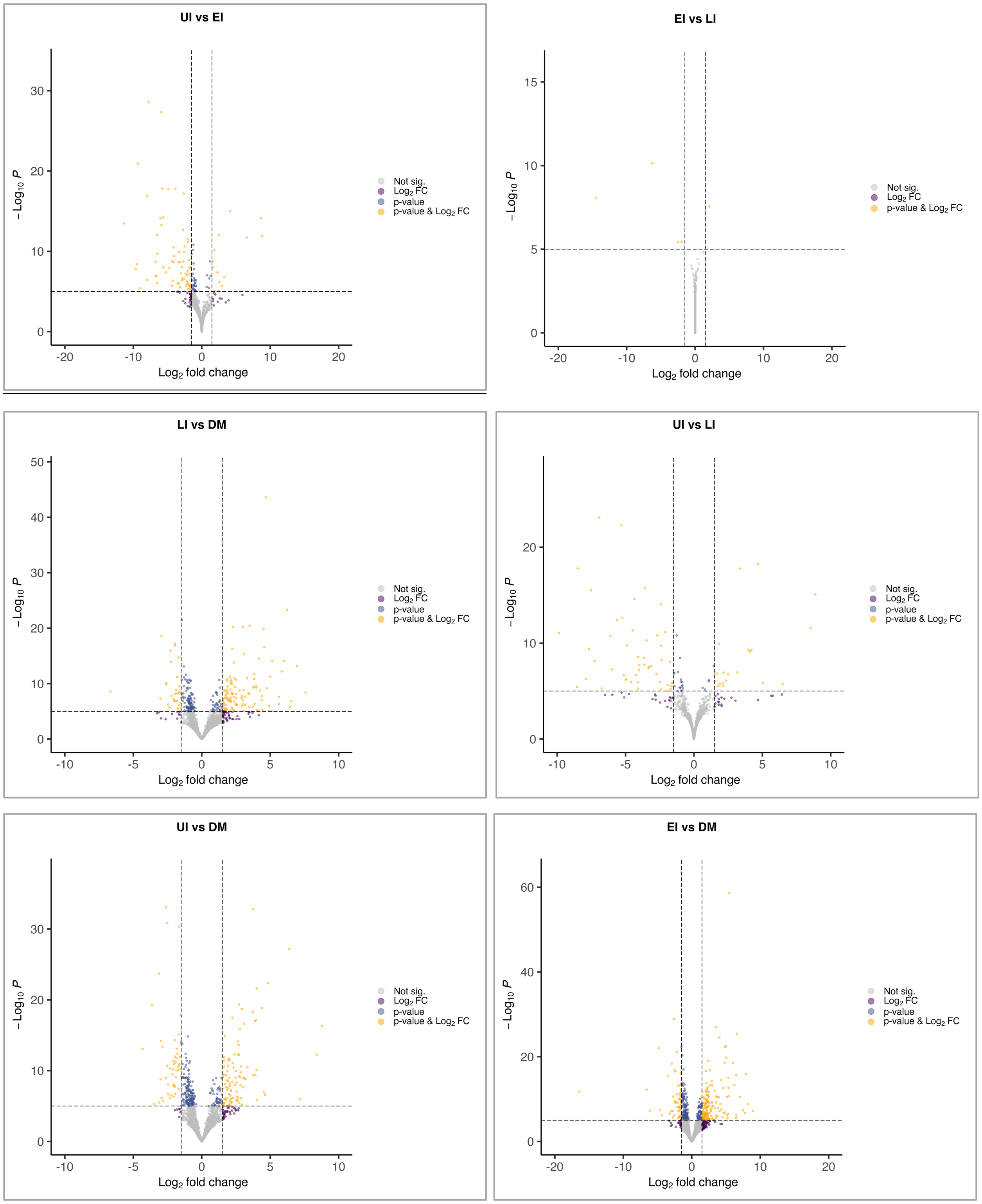
UI vs EI
EI vs LI
LI vs DM
UI vs LI
UI vs DM
EI vs DM
$-\text{Log}_{10}\,P$
$\text{Log}_2$ fold change
Not sig.
$\text{Log}_2$ FC
p-value
p-value & $\text{Log}_2$ FC

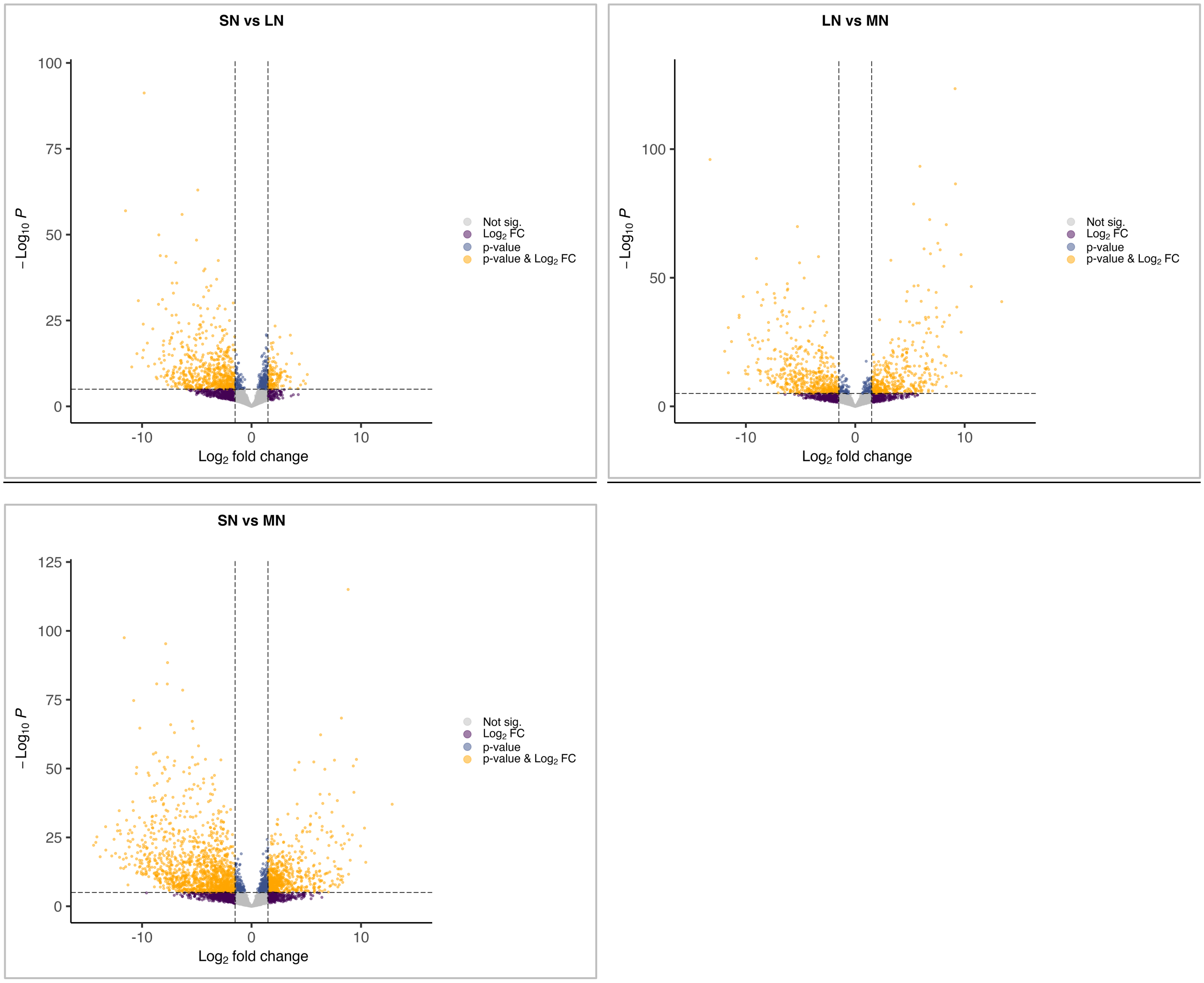


Supplementary Figure S1. Volcano plots of differentially expressed genes in pairwise comparisons. Plots comparing earwig samples are in black outline and those comparing nematodes are in red. Comparison groups are indicated on top of each plot. In earwig samples UI=uninfected, EI=early infection, LI=late infection, DM=during manipulation timepoints, similarly, SN=small nematodes, LN=large nematodes, and MN=manipulating nematodes. Yellow dots denote genes with FDR <0.05 and log2FC ≥2. Blue dots denote genes with FDR <0.05 and log2FC <2. Purple dots represent genes with FDR >0.05 and log2FC ≥2. Grey dots represent genes with FDR >0.05, log2FC <2. There are 9,007 earwig genes, and 9,722 nematode genes in their respective analysis.
